# Intrinsic neural timescales in the temporal lobe support an auditory processing hierarchy

**DOI:** 10.1101/2022.09.27.509695

**Authors:** Riccardo Cusinato, Sigurd L. Alnes, Ellen van Maren, Ida Boccalaro, Debora Ledergerber, Antoine Adamantidis, Lukas L. Imbach, Kaspar Schindler, Maxime O. Baud, Athina Tzovara

**Affiliations:** Institute of Computer Science, University of Bern, Switzerland; Center for Experimental Neurology - Sleep Wake Epilepsy Center - NeuroTec, Department of Neurology, Inselspital, Bern University Hospital, University of Bern, Switzerland; Swiss Epilepsy Center, Klinik Lengg, Zurich, Switzerland; Helen Wills Neuroscience Institute, University of California Berkeley, USA

## Abstract

During rest, intrinsic neural dynamics manifest at multiple timescales, which progressively increase along visual and somatosensory hierarchies. Theoretically, intrinsic timescales are thought to facilitate processing of external stimuli at multiple stages. However, direct links between timescales at rest and sensory processing, as well as translation to the auditory system are lacking. Here, we used intracranial electroencephalography in humans to show that in the auditory network, intrinsic neural timescales progressively increase, while the spectral slope flattens, from temporal to entorhinal cortex, hippocampus, and amygdala. Within the neocortex, intrinsic timescales exhibit spatial gradients that follow the temporal lobe anatomy. Crucially, intrinsic timescales at rest can explain the latency of auditory responses: as intrinsic timescales increase, so do the single-electrode response onset and peak latencies. Our results suggest that the human auditory network exhibits a repertoire of intrinsic neural dynamics, which manifest in cortical gradients with millimeter resolution and may provide a variety of temporal windows to support auditory processing.

## Introduction

The human brain gives rise to rich neural dynamics that mediate perception and cognition, and operate at multiple timescales (Honey et al., 2012; Murray et al., 2014). In the visual and somatosensory systems, intrinsic timescales manifest at rest, in ongoing neural activity: primary areas exhibit short timescales that may facilitate a quick reaction to incoming stimuli (Gao et al., 2020; Murray et al., 2014; Raut et al., 2020). These progressively increase while advancing through the cortical hierarchy, likely supporting integration of information (Chaudhuri et al., 2015; Murray et al., 2014). Whether a similar hierarchy of intrinsic dynamics exists in the temporal lobe, a hub for auditory processing, remains unknown.

In the auditory system, evidence for processing of external stimuli at multiple latencies stems from studying evoked responses. Primary auditory areas typically show fast and short-lasting responses, which may progressively increase while advancing to secondary areas (Camalier et al., 2012, Nourski et al., 2014). Beyond this ‘classical’ circuitry, an extensive network of adjacent regions is also sensitive to auditory input. These exhibit diverse response profiles and latencies, and include, for instance, the insula (Blenkmann et al., 2019), or the hippocampus and amygdala, which show slower, long-lasting responses to sounds (Halgren et al., 1980). Despite this diversity in auditory response profiles, a detailed characterization of temporal lobe dynamics at rest and their contribution to auditory processing remains an open question. In humans, in particular, a fine-grained measurement of neural dynamics in the temporal lobe can be challenging with non-invasive techniques (Tzovara et al., 2019), but evidence from invasive recordings remains limited.

Here, we hypothesized that intrinsic neural timescales at rest, estimated by characteristic latencies of the autocorrelation function (ACF) of intracranial electroencephalography (iEEG) signals, would show a hierarchical organization within an extended auditory network, which could, in turn, explain a hierarchy of neural responses to incoming auditory stimuli. We additionally hypothesized that non-oscillatory brain dynamics, characterized by the spectral exponent of aperiodic neural activity, which has been suggested to be a proxy of the excitation to inhibition balance (Gao et al., 2017), would also reveal a hierarchical organization across the temporal lobe.

## Results

We recorded iEEG in 270 electrodes from 11 patients with epilepsy, undergoing pre-surgical monitoring, presented with pure tones at random intervals (Figure 1A, Suppl. Table 1). Initially, we assessed a macroscopic organization of neural dynamics by dividing iEEG signals into regions of interest, selected based on the most consistent implantations across patients, targeting the entorhinal cortex (ENT), hippocampus (HIP), and amygdala (AMY) with their innermost electrodes, with additional electrodes covering the temporal and adjacent cortices (CTX) (Figure 1B, Supplemental Material for sub-divisions of cortical electrodes). Subsequently, we grouped all available electrodes together and assessed their spatial organization at a finer level, with respect to cortical and subcortical anatomies.

**Figure 1.**
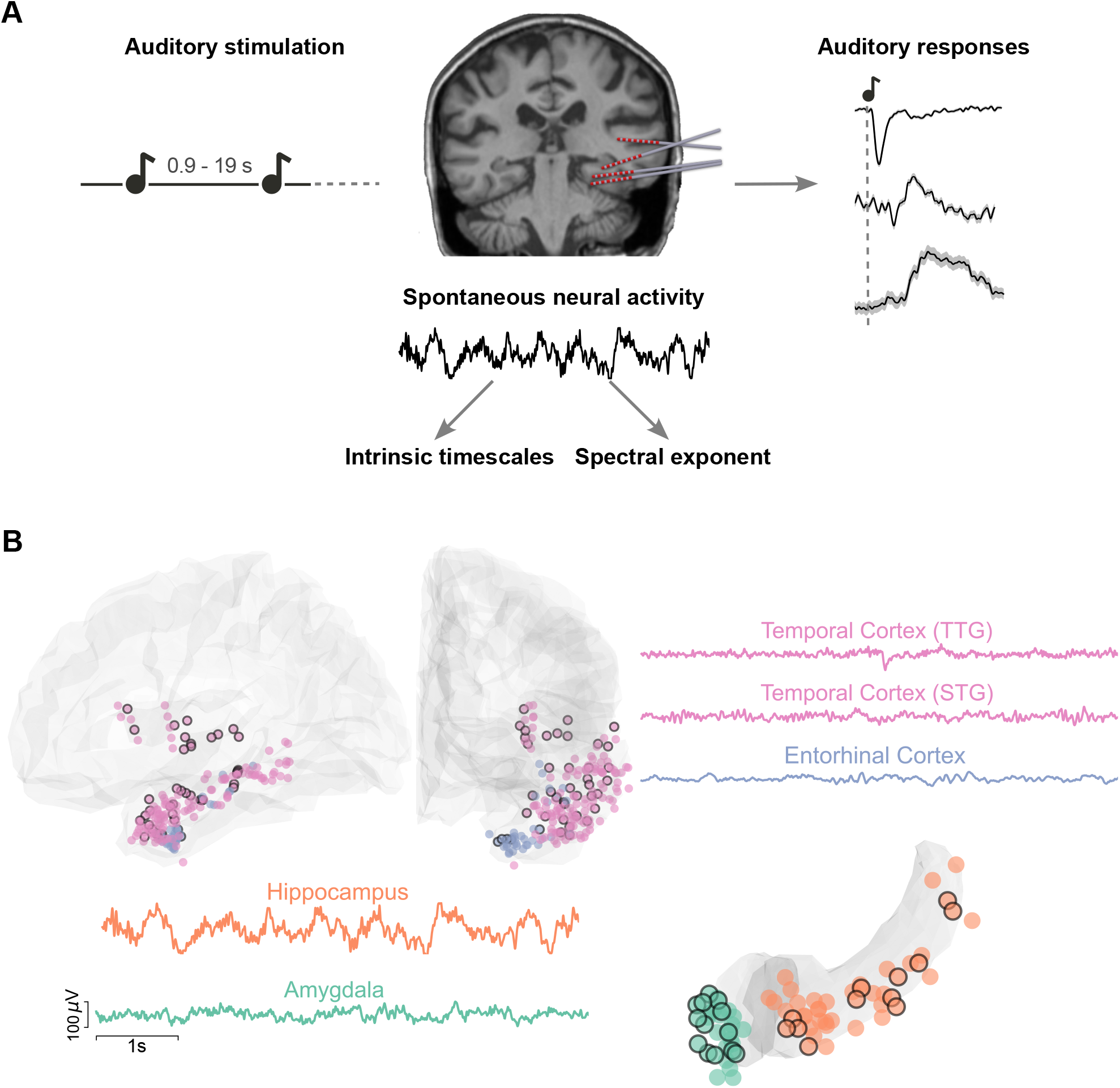
Experimental paradigm, electrode coverage, and exemplar iEEG traces. A. Summary of the main analyses and methodology. Left: schematic of the auditory stimulation protocol: Patients were presented with 100 ms pure tones, occurring at random intervals between 0.9-19 s. Middle: Example of implanted iEEG electrodes and exemplar raw trace of spontaneous neural activity from one electrode, before sound presentation, which is used to estimate intrinsic timescales and spectral exponents. Right: intracranial event-related potentials (iERPs) are esxtracted in response to the sounds. These are displayed for a schematic illustration of our protocol, for three exemplar electrodes, and are presented in more detail in Figure 3. B. Illustration of recorded electrodes (N=270) over the group of 11 patients. Black-circled electrodes are responsive to the auditory stimulation. As exemplar signals, we show iEEG traces from the transverse and superior temporal gyri (TTG and STG, pink), the entorhinal cortex (light blue), hippocampus (orange), and amygdala (green). Each of these regions exhibits characteristic and distinct dynamics at rest, displayed here over a 6 s segment.

We computed the autocorrelation function of baseline iEEG, which quantifies how similar a time series is to its past values across multiple time lags. The mean autocorrelation across brain regions shows a characteristic decay (Figure 2A) and for short time lags follows an ordering: electrodes in the temporal cortex have the most rapid decay, followed by electrodes in the entorhinal cortex, the hippocampus, and last, amygdala (Figure 2A, significant main effect of region for time-lags between 10 and 80 ms, p_corr_<0.05).

**Figure 2.**
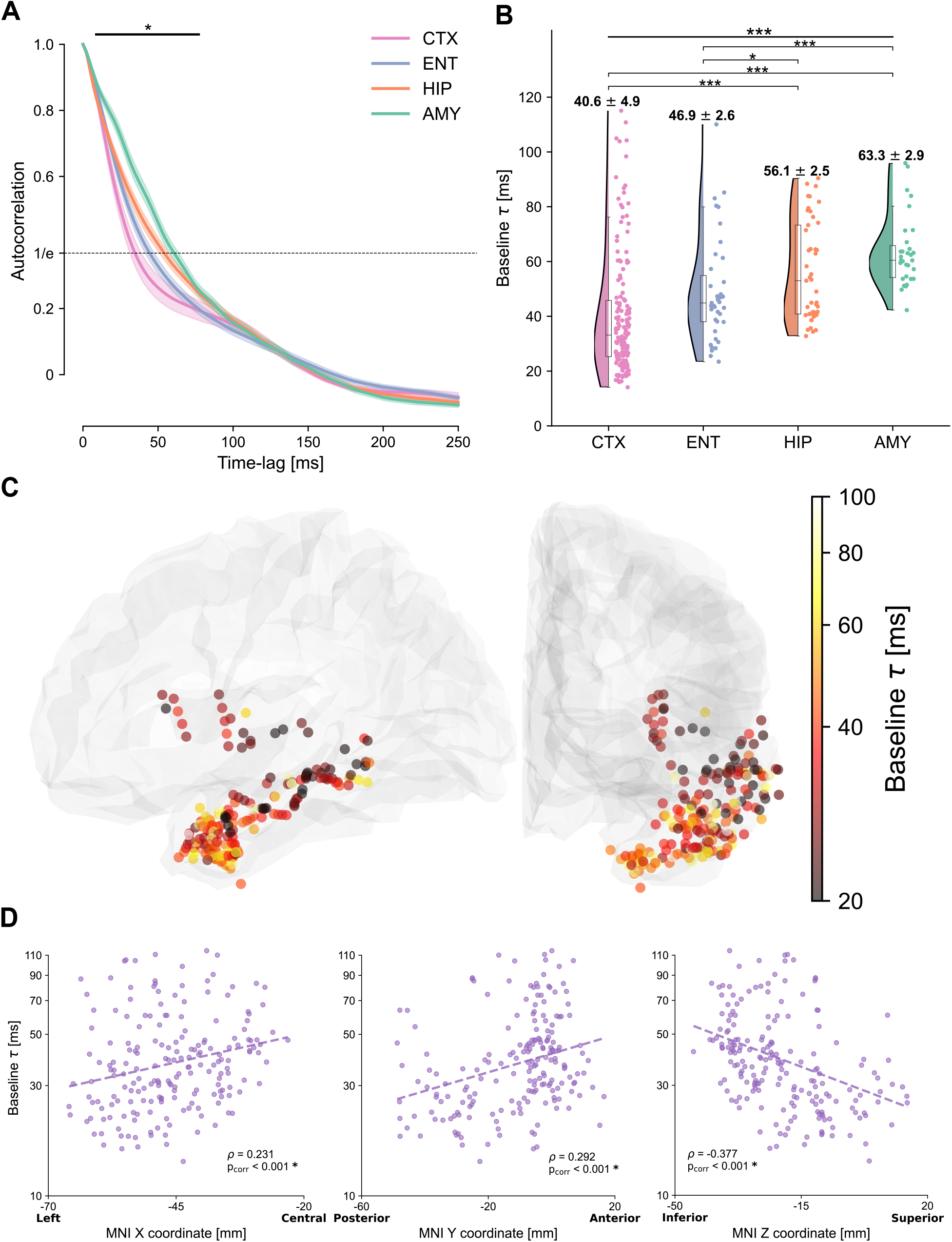
Autocorrelation function and intrinsic neural timescales at rest. A. Average autocorrelation function across electrodes and patients, for electrodes in the temporal (pink) and entorhinal (light blue) cortices, hippocampus (orange), and amygdala (green). The autocorrelation shows a significant main effect of region for time lags between 10 and 80 ms (horizontal solid bar). The dashed horizontal line at 1/e (inverse of natural logarithm) displays the value of the autocorrelation for which the characteristic timescales are extracted. B. Intrinsic timescales at baseline (τ), plotted for each electrode, show a main effect of region, with significantly faster timescales for the temporal and entorhinal cortices compared to the hippocampus and amygdala. C. The spatial organization of intrinsic timescales follows the cortical anatomy. Electrodes in the posterior/superior temporal cortex exhibit the fastest timescales, which progressively increase along the anterior/inferior axis. The color map quantifies the intrinsic timescale for each electrode on a logarithmic scale. For display purposes, all electrodes have been projected to the left hemisphere. D. Gradients of timescales spanning the cortex, plotted as timescales along the X, Y, and Z MNI coordinates of each electrode. Timescales significantly correlate with MNI coordinates in all three dimensions, tracking the cortical anatomy.

We then computed intrinsic neural timescales (τ), defined as the time lag at which the autocorrelation decays to a fixed threshold. Intrinsic timescales show a significant main effect of brain region (F(3,256)=27.313, p<10^−14^, mixed-effects model) (Figure 2B), and reveal a macroscopic hierarchy at rest: the temporal cortex exhibits significantly faster intrinsic timescales, compared to both hippocampus and amygdala (Suppl. Table 2). Within subregions of the cortex, intrinsic timescales tend to be slower in the pole, and faster in the transverse gyrus, while the superior, middle and inferior temporal cortex or the insula lie in between (Suppl. Figure 2-1). The entorhinal cortex is also significantly faster compared to other limbic areas, but not different from the temporal cortex (Suppl. Table 1).

At a finer scale, within the temporal and entorhinal cortices, intrinsic timescales at rest show a gradient that spans the temporal lobe through the postero-lateral (fast timescales) to the antero-medial (slow timescales) axis (Figure 2C). This gradient is particularly prominent in the Y and Z directions that primarily define the temporal lobe orientation (Figure 2D, ρX=0.231, p_X_<10^−5^; ρ_Y_=0.292, p_Y_<10^−8^; ρ_Z_=-0.377, p_Z_<10^−11^, accounting for inter-subject variability and Bonferroni corrected). The spatial distribution of timescales in the hippocampus and amygdala is by contrast less defined (Suppl. Figure 2-2).

We next investigated whether intrinsic timescales at rest could explain the timing of auditory processing. iERPs across brain regions show striking qualitative differences in terms of amplitude and response timing (Figure 3A). To quantify these diverse auditory response profiles, we computed the response onset and peak latencies of electrodes showing a significant 1-40 Hz auditory response (N=67 out of 270 total electrodes, Figure 3A/B). Cortical electrodes generally show faster responses than hippocampal and amygdalar ones both for onset and peak, but at the group level, there is no significant effect of region (F(3,55)=1.867, p=0.146; F(3,55)=2.774, p=0.0499 for onset/peak, Figure 3B and Suppl. Figure 3-1 for sub-regions).

**Figure 3.**
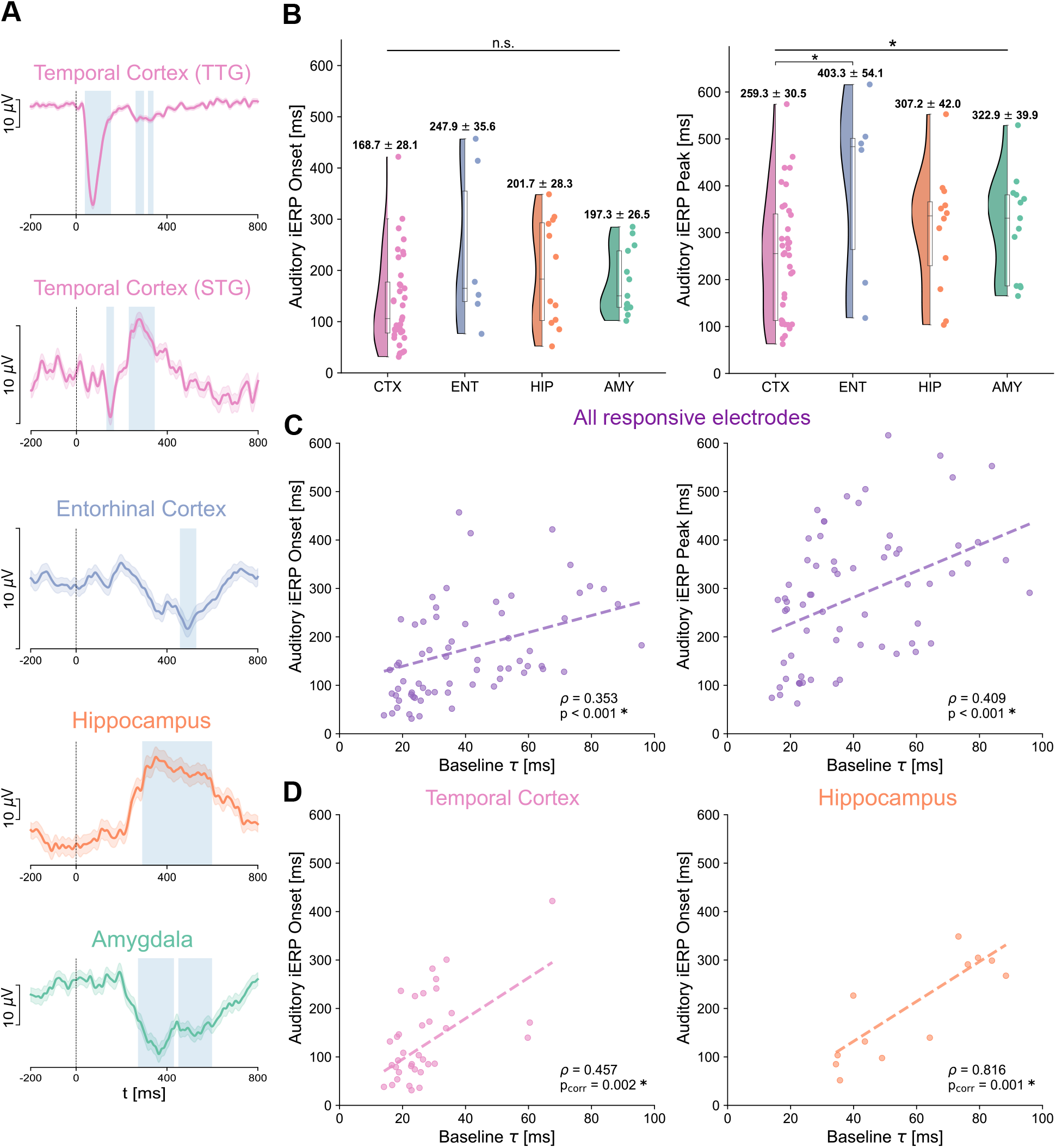
Onset and peak latencies of auditory responses across brain regions and their relation to intrinsic timescales at rest. A. Exemplar auditory responses for each of the recorded regions (1-40 Hz iERPs). Time 0 corresponds to sound onset. Auditory responses in the transverse temporal gyrus (TTG) are the earliest, shorter-lasting, and exhibit the largest amplitude (top plot). Responses in other cortical regions, e.g. the superior temporal gyrus (STG), have a relatively early onset, and later peak, while responses in the entorhinal cortex, hippocampus, and amygdala (third to fifth row) are typically smoother, long-lasting, and with later peaks. The variability in response amplitudes is indicated by the different spans of a 10μV scale on the y-axis. B. Auditory response onsets (left panel) and peaks (right panel) for all responsive electrodes. The temporal cortex shows the earliest onset and peak across all brain regions, with responses starting on average at 168.7 ms, and peaking at 259 ms post sound onset, followed by the hippocampus/amygdala, and entorhinal cortex. C. Regression of intrinsic timescales τ at baseline (x-axis) on auditory iERP onsets (y-axis, left panel) and peaks (y-axis, right panel) across all responsive electrodes. Regression of intrinsic timescales on both onsets and peaks is highly significant, accounting for across-patient variations, suggesting that intrinsic timescales can explain the timing of auditory responses at the single electrode level. D. A significant regression of characteristic timescales on iERP onsets also persists within the temporal cortex (left panel), and hippocampus only (right panel), but not in the amygdala or entorhinal cortex (Suppl. Figure 3-2).

This variability in onset and peak latencies within and between brain regions can be partially explained when accounting for differences in intrinsic timescales (Figure 3C). We computed a regression of intrinsic timescales on response latencies, which shows a highly significant main effect of timescale at baseline both on response onset (ρ=0.353, p=0.0009) and peak latencies (ρ=0.409, p=0.0005). The significant regression on onset latency holds for electrodes within the temporal cortex (ρ=0.457, p_corr_=0.0017) and hippocampus (ρ=0.816, p_corr_=0.0013, Figure 3D) (Suppl Figure 3-2 for other sub-regions). These results suggest that regions characterized by fast intrinsic timescales exhibit a fast reaction to incoming auditory stimuli, while the hippocampus, amygdala, and entorhinal cortex are mediated by slower ongoing dynamics and slower auditory responses.

To confirm the observed hierarchy of intrinsic neural dynamics, we additionally characterized the non-oscillatory part of the power spectra by their spectral exponent (slope in log-log space) in the ranges 20-35 and 80-150 Hz (Figure 4A). Electrodes in the temporal cortex have the steepest 20-35 Hz exponent, followed by the entorhinal cortex, hippocampus, and amygdala (Figure 4B, significant effect of region, F(3,256)=80.665, p<10^−16^), confirming the ordering observed for intrinsic timescales (Suppl. Table 3). The spectral exponent in the 80-150 Hz range also shows a significant main effect of region (F(3,256)=79.156, p<10^−16^, Figure 4C), mainly driven by the difference between the hippocampus and all other regions (Suppl. Table 4). Similar to the intrinsic timescales, the 20-35 Hz spectral exponent shows a significant, albeit weaker, correlation with MNI X and Z coordinates (Suppl Figure 4-3 and Suppl Figure 4-4 for within hippocampus/amygdala), providing further support for a gradient organization of neural dynamics within the extended auditory cortical network.

**Figure 4.**
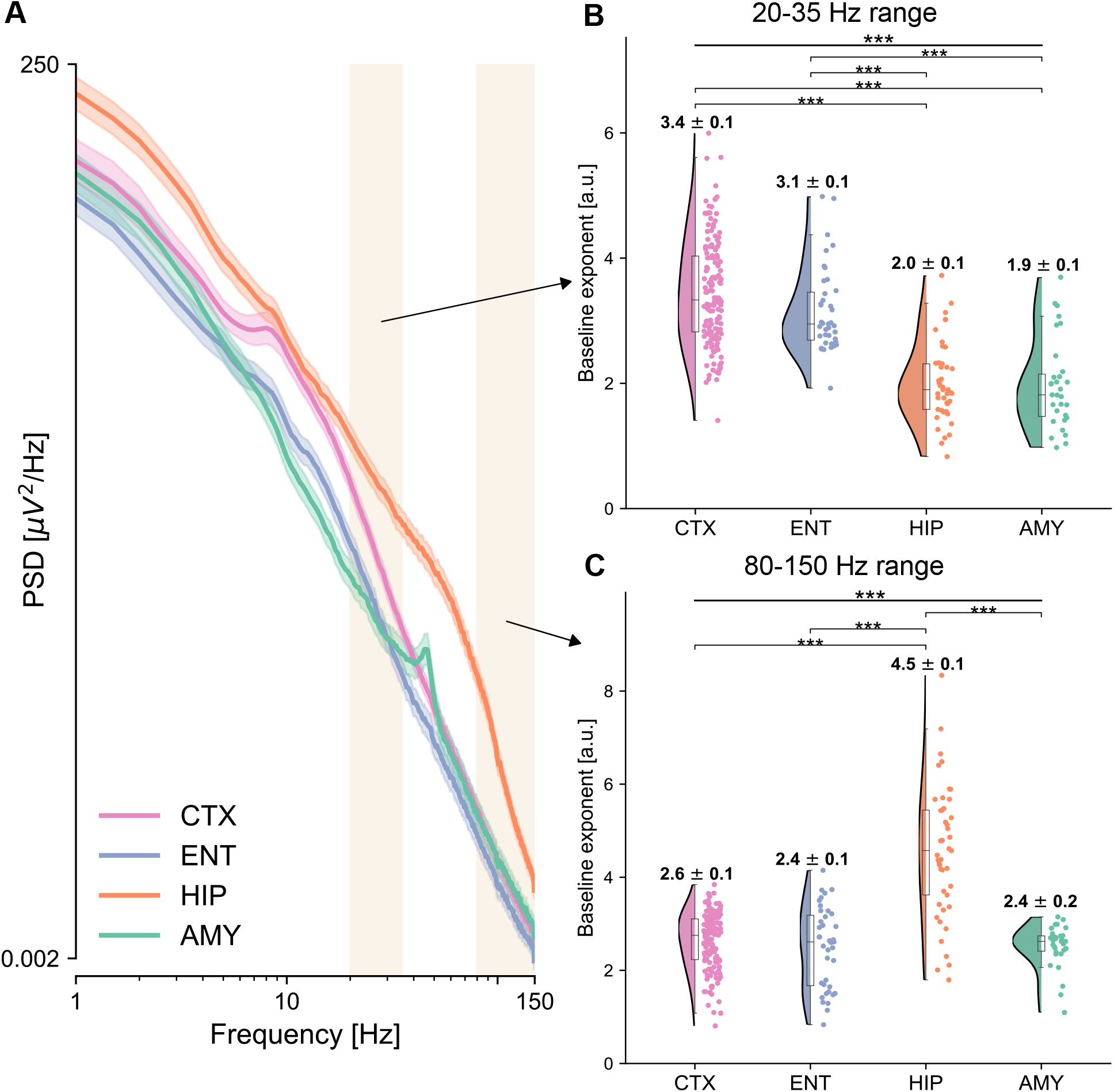
Power spectra and spectral exponents across brain regions. A. Average power spectra are displayed for the four regions of interest. Cortex (pink) exhibits a characteristic oscillatory peak around 10 Hz, and a relatively fast decay, while the hippocampus (orange) displays the strongest power, which for low frequencies decays relatively gently, but after 70 Hz much faster. The shaded rectangles highlight the two frequency ranges for which the spectral exponent is computed, at 20-35 Hz, and at 80-150 Hz. x- and y-axes are plotted in logarithmic scales. B/C. Spectral exponent at 20-35 Hz (B) and 80-150 Hz (C), for each electrode and region of interest. The spectral exponent in the 20-35 Hz range shows a significant main effect of region, with the temporal cortex having the steepest exponent followed by the entorhinal cortex, the hippocampus, and amygdala, which have flatter exponents. The 20-35 Hz exponent values for cortical subdivisions can be found in Suppl Figure 4-1. The spectral exponent at 80-150 Hz also shows a significant effect of region, with the hippocampus having the steepest exponent among all other regions, compatible with the “knee” observed in the average power spectra (panel A). The corresponding offsets for both ranges are reported in Suppl Figure 4-2.

## Discussion

We provide evidence for a hierarchy of intrinsic neural dynamics in the extended human auditory network at rest, which in turn explains a hierarchy in the processing of incoming auditory stimuli. The temporal cortex assumes a “low” position along this hierarchy, highlighted by short intrinsic timescales, which may mediate short temporal receptive windows (Hasson et al., 2008), and by a steep 20-35 Hz spectral exponent, which may indicate a shift towards inhibitory activity (Gao et al., 2017). On the contrary, the hippocampus and amygdala exhibit longer intrinsic timescales and a flatter exponent, likely indicative of an integrative function (Golesorkhi et al., 2021; Murray et al., 2014). Our findings are in line with previous reports of a hierarchical organization of intrinsic timescales in the visual and somatosensory modalities, which progressively increase along the cortical hierarchy (Murray et al., 2014), and also with reports of a hierarchical organization in synaptic excitation (Wang, 2020). Although here we were not able to directly measure excitation/inhibition, the steepness of the spectral exponent around the 20-35 Hz range has been suggested to reflect the excitation to inhibition balance (Gao et al., 2017), or levels of neural “noise” (Alnes et al., 2021). The particularly steep hippocampal spectral exponent for high frequencies, by contrast, reflects the abrupt change of slope in the hippocampal power spectrum, which forms a “knee” at around 70 Hz (Figure 4A).

We additionally show that the hierarchy of intrinsic neural dynamics of the extended auditory network manifests as a continuous gradient along the anatomy of the temporal lobe, both for intrinsic timescales and spectral exponents. We did not find evidence for gradients of timescales within the hippocampus and amygdala, contrary to a recent magnetic resonance imaging study (Raut et al., 2020), possibly because of a sparser coverage or because these may only manifest at very slow dynamics.

Although several studies have posited that short intrinsic timescales may mediate fast responses to incoming stimuli, we now provide direct evidence for the auditory modality. We show, in the same patients and recordings, that the diversity of intrinsic timescales partially explains the richness of auditory response onset and peak latencies. Although our analyses are correlational, we posit that this repertoire of intrinsic timescales at rest may support the auditory process itself, providing a variety of processing windows (Golesorkhi, et al., 2021). How this hierarchy supports processing of more complex stimuli, or whether it expands to structures of the midbrain can be the topic of future investigations.

In summary, our results show a hierarchy of neural dynamics in the extended human auditory network that manifests across cortical and limbic structures, exhibits anatomical gradients with millimeter resolution and can explain the temporal richness of neural responses to auditory stimuli.

## Methods

### Patients

We recorded intracranial EEG in 11 neurosurgical patients (4 women, median age=32 years, min=27, max=56) with drug-refractory epilepsy who had been implanted with depth electrodes to identify seizure foci (Supplemental Table 1 for a detailed patient description). Electrode locations were based on clinical criteria only. Recordings took place at the EPI Clinic, Zurich, and at the Inselspital, Bern. Patients provided written informed consent prior to participation in this research study, approved by institutional ethics review boards of the Canton of Zurich (PB-2016-02055), and Inselspital, Bern (# 2018-01387). All experiments were performed in accordance with the 6^th^ Declaration of Helsinki.

### Experimental protocol

Patients were presented with auditory stimuli consisting of pure tones at three frequencies (500, 1250, 2500 Hz) with a random interstimulus interval between 0.9 and 19 seconds. Each tone had a duration of 100 ms with 5 ms on/off ramps to avoid clicks. Interstimulus interval and tone frequency were drawn from a pseudorandom distribution such that each was played 120 times per hour (in total 360 tones per hour). Auditory stimuli were presented via in-ear headphones, and their intensity was adjusted individually for each patient at a comfortable level. Patients were instructed to relax and ignore the sounds. Some of the patients were additionally presented with the auditory stimuli during sleep, at a later session, which was not analyzed in the context of the present study.

### iEEG recordings & preprocessing

Depth electrodes were used for iEEG recordings (DIXI Medical, 3 patients; Ad-Tech Medical, 8 patients) targeting different brain regions and varying from eight to eighteen platinum iEEG contacts along their shaft. Data were recorded at 4096 or 1024 Hz. Recordings with 4096 Hz sampling rate were downsampled offline to 1024 Hz. All data were visually inspected to exclude electrodes with persistent spiking activity. Continuous data were notch filtered around 50 Hz and harmonics, and re-referenced with a bipolar scheme, i.e. each electrode to the closest one in the same electrode lead outwardly, to remove any source of widespread noise. This was done to retain a local signal and mitigate effects of volume conductance, following recommendations in the analysis of iEEG data (Lachaux et al., 2012; Mercier et al., 2022). Peri-stimulus epochs were then extracted, spanning from -5 s before the sounds’ onset to 5 s post-stimulus onset. Only epochs that did not overlap with another sound in this period were kept. All epochs were then visually inspected and any epochs with remaining artifacts were rejected. The baseline period of each epoch was defined as the interval [-1,0] s preceding the sounds. For studying auditory responses (see Responsive electrodes section), the raw signal from all electrodes was additionally band-pass filtered between 1-40 Hz. Processing of iEEG data was performed using MNE python (Gramfort et al., 2013).

### Electrode localization

Electrodes were localized on post-implant computed tomography (CT) scans using the Lead-DBS toolbox (Horn & Kühn, 2015) and transformed into standard MNI coordinates for group analyses. The post-implant CT scan was registered to a pre-implant structural T1-weighted magnetic resonance imaging (MRI) scan from which anatomical labels were reconstructed using the FreeSurfer toolbox and the Destrieux atlas. Subsequently, electrode coordinates identified on the post-implant CT scans were mapped to their corresponding anatomical regions identified on the pre-implant MRI. Anatomical label assignment was validated for all electrodes by an expert neurologist, who verified their location and additionally ensured that none of the electrodes that were included in our analyses were in white matter. The available electrodes were divided across four regions of interest, covering the temporal cortex, the insula due to its prominent auditory responses, entorhinal cortex, hippocampus, and amygdala. This resulted in N=270 electrodes in total, with a median=25, min=8 and max = 37 electrodes per patient.

### Intrinsic neural timescales

For estimating intrinsic neural timescales, we first computed the Autocorrelation function (ACF) on each epoch during 1 s baseline period (function *acf* from Python’s statsmodels (Seabold & Perktold, 2010)). The resulting ACFs across epochs were then averaged to yield a single ACF for each electrode. Following previous literature (Chaudhuri et al., 2015; Gao et al., 2020; Golesorkhi, et al., 2021; Murray et al., 2014), we then defined the “intrinsic timescale” of each electrode as the time lag at which the ACF reaches the value 1/e, consistent with an analytical decay of the form *f(t)=exp(-t/τ)*. The precise time-lag was computed by interpolating with a spline fit to the ACF, as in (Raut et al., 2020).

### Power spectral density and spectral exponent

For estimating the spectral exponent, we computed power spectra with a Hann-windowed and detrended Fourier transform on the baseline period (function *spectrogram* from Python’s scipy (Virtanen et al., 2020)). Power spectra were averaged using a “meanlog” approach, i.e. taking the mean of the logarithm of the power spectrum across epochs, to yield a single power spectrum density for each electrode.

The spectral exponent was then computed on each electrode’s average power spectrum density using the standard implementation of the spectral parameterization algorithm (Donoghue et al., 2020) in the “fixed” mode (linear fit in log-log plot) in two different frequency ranges: a lower one, at 20-35 Hz, and a higher one, at 80-150 Hz. The lower range was chosen following a large body of literature in order to avoid low-frequency knees, high-power peaks and spectral plateaus, and has been previously linked to excitation to inhibition balance (Gao et al., 2017; Gerster et al., 2021; Lendner et al., 2020). The higher range was chosen as a typical high frequency range that is often computed in iEEG studies, as a proxy to neuronal firing (Lachaux et al., 2012). The spectral exponent was computed as the slope of non periodic parts of the power spectra observed at each electrode via a standarized approach in the FOOOF package (Donoghue et al., 2020) (parameters for the fitting: peak_threshold = 2, min_peak_height: 0.1, peak_width_limits: [1, 10], with max_n_peaks=2 for the lower range and 0 for the higher one). Fits for every electrode were visually inspected, and any electrodes with clear artifacts on the power spectra, or where the fit was particularly noisy were excluded to ensure an accurate estimation of the spectral exponent. After this step, all remaining electrodes (N= 270) had R^2 fits of at least 0.8. For our main analyses we were interested in spectral exponents, reported in the main text. The corresponding offsets are reported in Suppl Figure 4-2, for reasons of completeness. Amygdalar electrodes from two patients had a prominent peak in their power spectra around 40 Hz (Figure 4A), which was found for electrodes of the amygdala only, and not other electrodes, and to the best of our knowledge was unrelated to any sources of noise, or pathological findings in these patients. We confirmed that fitting of the spectral exponent was not affected by these peaks in none of the two patients, which were outside the range of our fits.

### Responsive electrodes

Responsive electrodes were identified following common approaches in the field of iEEG (Dürschmid et al., 2016). Briefly, differences between the average signal in post-stimulus time points, *Ā*(*t*), and over the entire baseline, 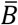, were compared with surrogate distributions computed by random-shifting of the original epochs for i=1,…,1000 iterations ({*A*_*i*_*(t)-B*_*i*_}_*i=1…1000*_). Response time points were considered significantly different from the baseline if 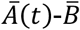 fell outside the outer 5% interval of the permuted distribution. Additionally, only electrodes with at least one consecutive response lasting more than 50 ms were kept, to correct for multiple comparisons, as commonly done in the field (Guthrie & Buchwald, 1991; Haller et al., 2018; Kam et al., 2021). The post-stimulus time-points were restricted to the interval [10, 600] ms, to control for too early and too late onsets that would be biologically implausible. We defined the peak latency as the time between the sound onset and the maximum absolute difference voltage from baseline.

### Statistical analyses

Statistical tests have been conducted in R version 4.2.0 (R Development Core Team, 2020) using Linear Mixed-Effects models (LMEs) with a random intercept term corresponding to the patient, to account for the fact that electrodes were recorded from different patients (Yu et al., 2022) (implemented with *nlme* package (Lindstrom & Bates, 1990)). The omnibus tests for the “brain region” factor were computed with F-tests, while post-hoc pairwise comparisons were computed with Tukey’s range test, controlling for multiple comparisons (implemented with *emmeans* package). In the case of omnibus tests on multiple time lags (Fig. 2a) and tests over multiple MNI coordinates, p-values have been Bonferroni-corrected. For regression analyses, we used LMEs with a continuous predictor and a random intercept accounting for across-patient variability. We computed correlation values starting from R^2^ as described in (Nakagawa & Schielzeth, 2013) and took the square root, mimicking a fixed-effects-only linear model (implemented with *MuMIn* package (Kamil Barton, 2020)). P-values were computed with F-tests, correcting with Bonferroni when regressing on each level of the region factor separately (p_corr_).

## Acknowledgments

This work is supported by the Inselspital University Hospital Bern, the Interfaculty Research Cooperation “Decoding Sleep: From Neurons to Health & Mind” of the University of Bern (AA, IB, AT, SLA), the Swiss National Science Foundation (AA, #320030_188737 AT), the European Research Council (CoG-725850 to A.A.), the Synapsis Foundation (AA), the University of Bern (AA), and the Fondation Pierre Mercier pour la science (AT). The authors thank Prof. Johannes Sarnthein for support with recordings and feedback on the manuscript.

## Data and code availability

Because of the sensitive nature of the data, data and code can be made available from the corresponding author upon reasonable request.

## Supplementary material

### Supplementary tables

**Supplementary Table 1.** Overview of patients dataset. We collected a total of 270 electrodes from 11 patients, with a median of 25 electrodes per patient and minimum and maximum of 8 and 37 electrodes. For each patient, we report the hospital where the data were collected, the # of electrodes used for our analyses, the hemisphere(s) where the electrodes were implanted and the regions sampled from the retained electrodes.

| Patient ID | Clinic | # of electrodes analyzed | Hemisphere | Regions |
| --- | --- | --- | --- | --- |
| 1 | Zürich | 25 | L + R | CTX, ENT, HIP, AMY |
| 2 | Bern | 17 | R | CTX, ENT, HIP |
| 3 | Zürich | 34 | L + R | CTX, ENT, HIP, AMY |
| 4 | Zürich | 30 | L + R | CTX, ENT, HIP, AMY |
| 5 | Zürich | 28 | L + R | CTX, ENT, HIP, AMY |
| 6 | Zürich | 20 | L | CTX, ENT, HIP, AMY |
| 7 | Zürich | 37 | L + R | CTX, ENT, HIP, AMY |
| 8 | Bern | 24 | L | CTX, ENT, HIP |
| 9 | Zürich | 28 | L + R | CTX, ENT, HIP, AMY |
| 10 | Zürich | 19 | R | CTX, ENT, HIP, AMY |
| 11 | Bern | 8 | L | CTX, HIP |

**Supplementary Table 2.** Pairwise comparisons of intrinsic neural timescales across regions of interest. The first column lists each of the six pairwise comparisons, the second one the t-values of the post-hoc t-test, and the third column the related p-values. All pairs of cortical-limbic areas have significant differences in their intrinsic timescales, while the differences between temporal/entorhinal cortex and hippocampus/amygdala are non-significant. The timescale values are computed through a mixed-effects model with a patient-specific random effect and p-values are corrected for multiple comparisons via the Tuckey range test.

| Comparison | t-value | p-value |
| --- | --- | --- |
| CTX-ENT | -2.383 | 0.083 |
| CTX-HIP | -6.099 | $2.34 \times 10^{-8}$ |
| CTX-AMY | -7.716 | $1.69 \times 10^{-12}$ |
| ENT-HIP | -2.817 | 0.027 |
| ENT-AMY | -4.635 | $3.36 \times 10^{-5}$ |
| HIP-AMY | -2.067 | 0.167 |

**Supplementary Table 3.**
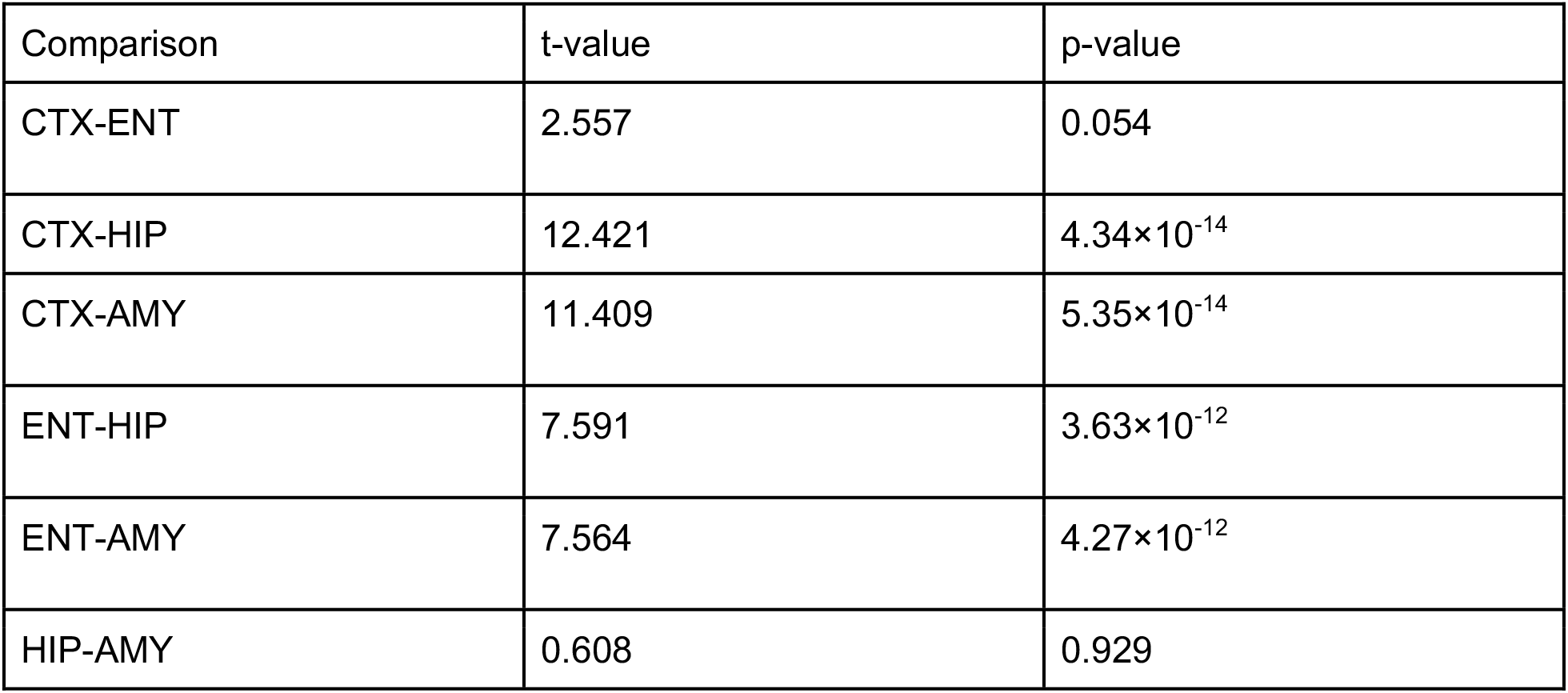
Pairwise comparisons of 20-35 Hz spectral exponents among the four regions of interest. The first column lists each of the six pairwise comparisons, the second one the t-values of the post-hoc t-test, and the last one the related p-values. All pairs of cortical-limbic areas have significant differences in their 20-35 Hz exponent, while the difference between temporal and entorhinal cortex is slightly above significance threshold. The spectral exponent values are computed through a mixed-effects model with a patient-specific random effect and p-values are corrected for multiple comparisons via the Tuckey range test.

| Comparison | t-value | p-value |
| --- | --- | --- |
| CTX-ENT | 2.557 | 0.054 |
| CTX-HIP | 12.421 | $4.34 \times 10^{-14}$ |
| CTX-AMY | 11.409 | $5.35 \times 10^{-14}$ |
| ENT-HIP | 7.591 | $3.63 \times 10^{-12}$ |
| ENT-AMY | 7.564 | $4.27 \times 10^{-12}$ |
| HIP-AMY | 0.608 | 0.929 |

**Supplementary Table 4.** Pairwise comparisons of 80-150 Hz spectral exponents among the four regions of interest. The first column lists each of the six pairwise comparisons, the second one the t-values of the post-hoc t-tests, and the last one the related p-values. Only the comparisons between hippocampus and the other areas are significant due to the very steep slope of hippocampal electrodes in the high-gamma range. The spectral exponent values are computed through a mixed-effects model with a patient-specific random effect and p-values are corrected for multiple comparisons via the Tuckey range test.

| Comparison | t-value | p-value |
| --- | --- | --- |
| CTX-ENT | 1.551 | 0.408 |
| CTX-HIP | -14.214 | $4.31 \times 10^{-14}$ |
| CTX-AMY | 1.631 | 0.363 |
| ENT-HIP | -12.321 | $4.35 \times 10^{-14}$ |
| ENT-AMY | 0.195 | 0.997 |
| HIP-AMY | 11.650 | $4.90 \times 10^{-14}$ |

### Supplementary figures

**Supplementary Figure 2-1.**
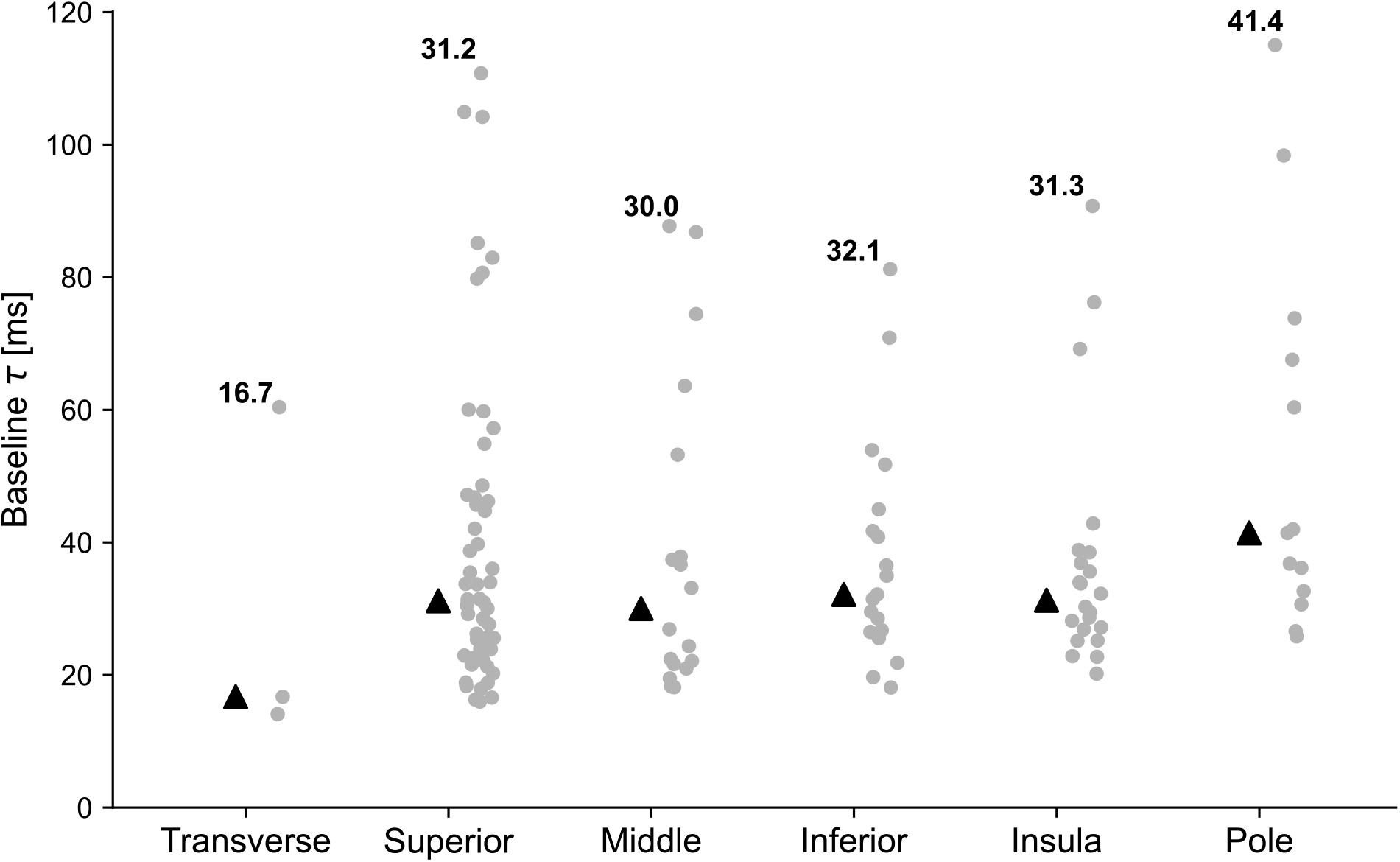
Intrinsic neural timescales across cortical sub-divisions. Intrinsic neural timescales of cortical electrodes, divided in anatomical subregions. Black triangles indicate median values, while the actual value is written for each region. The fastest timescales are observed for the transverse temporal gyrus, at 16.7 ms. All of the remaining cortical sub-divisions apart from the temporal pole have median timescales around 30-32 ms, with variations within them. The temporal pole by contrast has the longest timescales, with a median of 41.4 ms. The anatomical location of each electrode can be found in Figure 2 of the main text. Transverse: transverse temporal gyrus; Superior: superior temporal gyrus and sulcus; Middle: middle temporal gyrus; Inferior: inferior temporal gyrus and sulcus; Insula: inferior circular sulcus of insula; Pole: temporal pole.

**Supplementary Figure 2-2.**
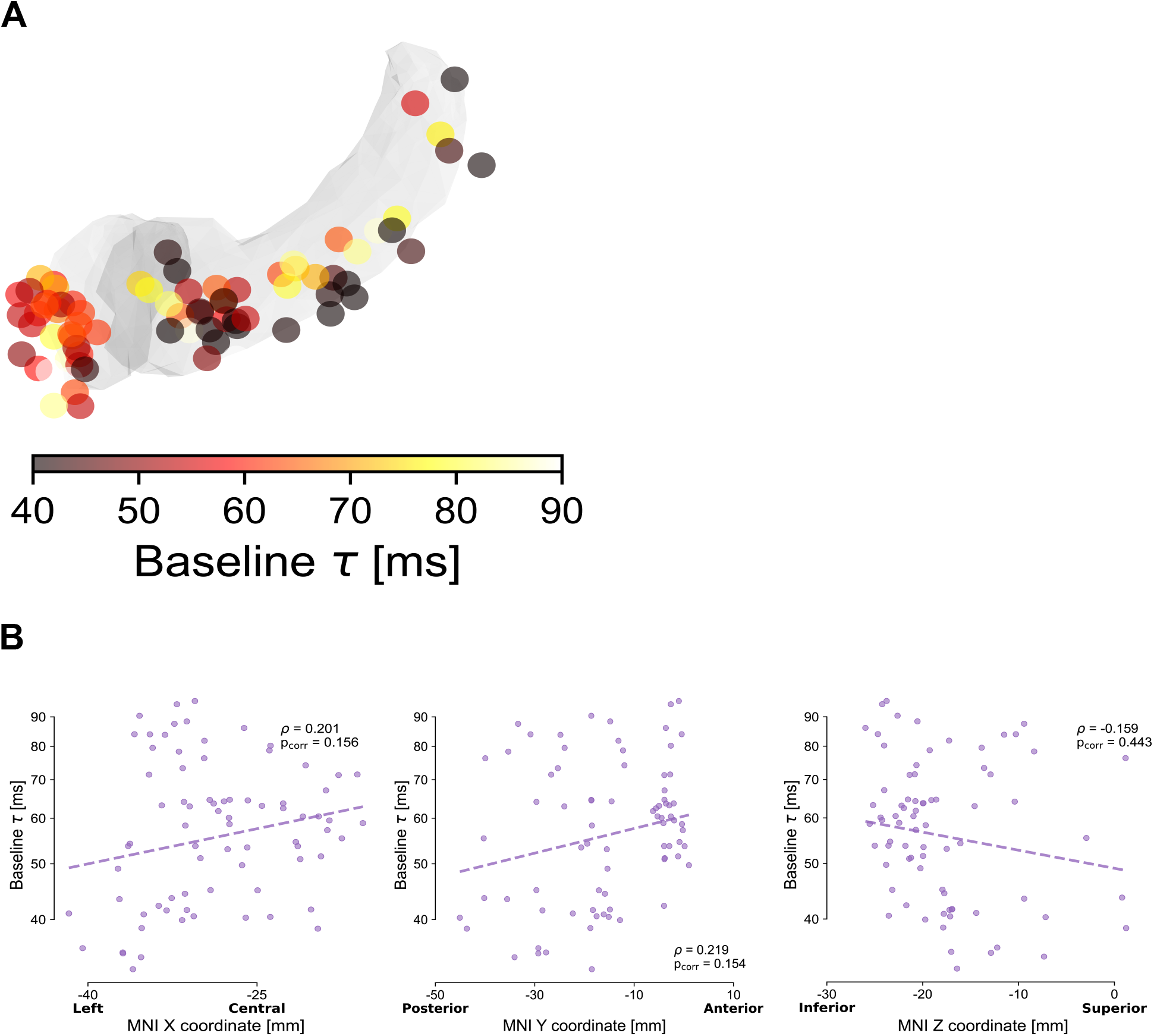
Intrinsic neural timescales at rest in the hippocampus and amygdala. A. Anatomical organization of intrinsic timescales throughout the hippocampus and amygdala, with the color map quantifying the intrinsic timescale for each electrode. B. Correlations between MNI coordinates and intrinsic timescale (τ) across electrodes. Although τ tends to be slower for anterior electrodes, and in particular for the amygdala, correlations in the X, Y, and Z directions are not significant (ρ_X_=0.201, p_X_=0.156; ρ_Y_=0.219, p_Y_=0.154; ρ_Z_=-0.159, p_Z_=0.443; p-values after Bonferroni correction). For display purposes, all electrodes have been projected to the left hemisphere.

**Supplementary Figure 3-1.**
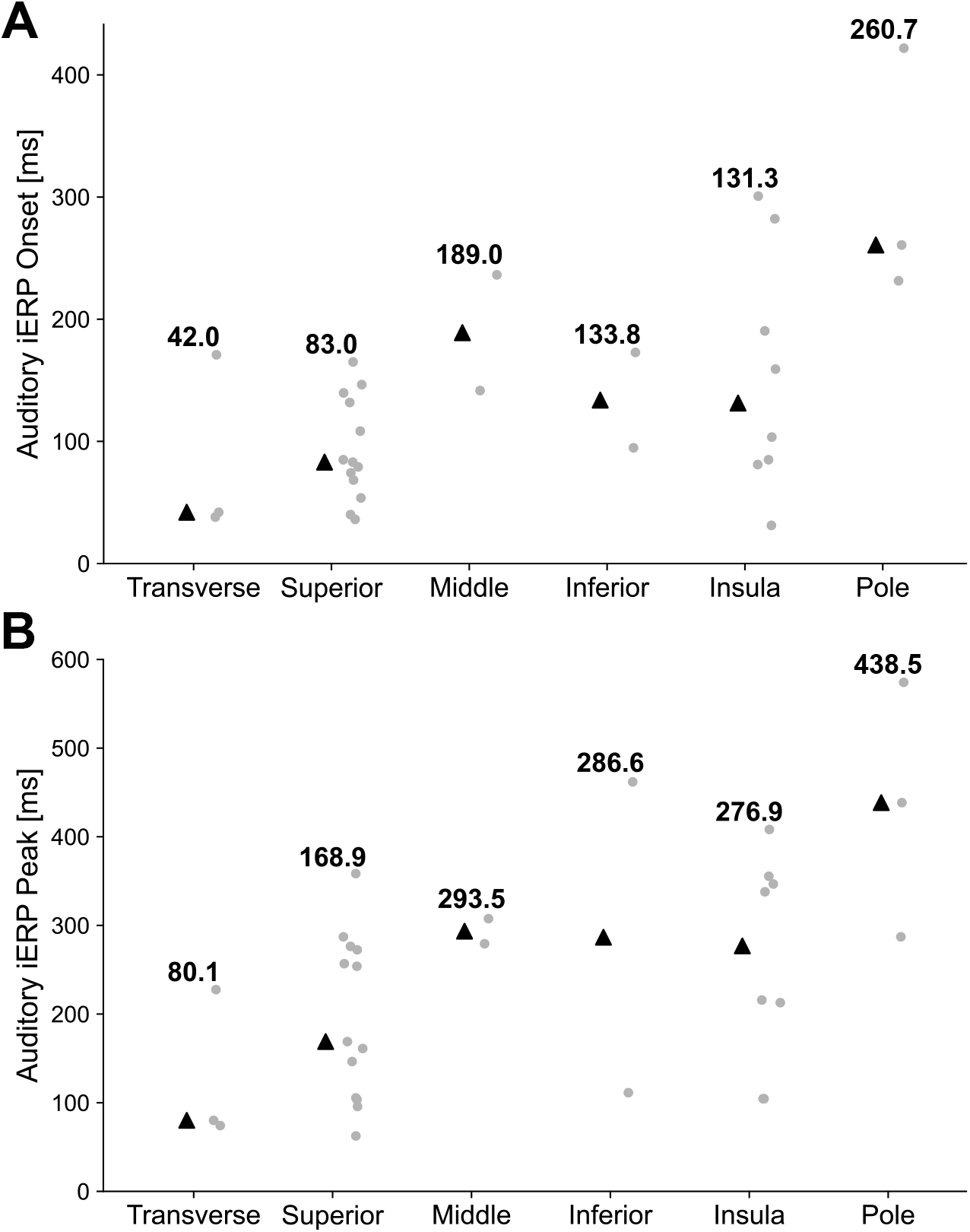
Auditory response latencies across cortical sub-regions. Onset latency (A) and peak latency (B) of Temporal cortex electrodes, divided by anatomical subregion. Black triangles indicate median values, with the actual value written for each sub-region. The transverse temporal gyrus has the earliest onsets (median at 42.0 ms) and peaks (median at 80.1 ms), followed by the superior temporal gyrus/sulcus, at 83 ms for onset and 168.9 ms for peak. The latest responses are observed for the temporal pole, with a median onset at 260.7 ms. Transverse: transverse temporal gyrus; Superior: superior temporal gyrus and sulcus; Middle: middle temporal gyrus; Inferior: inferior temporal gyrus and sulcus; Insula: inferior circular sulcus of insula; Pole: temporal pole.

**Supplementary Figure 3-2.**
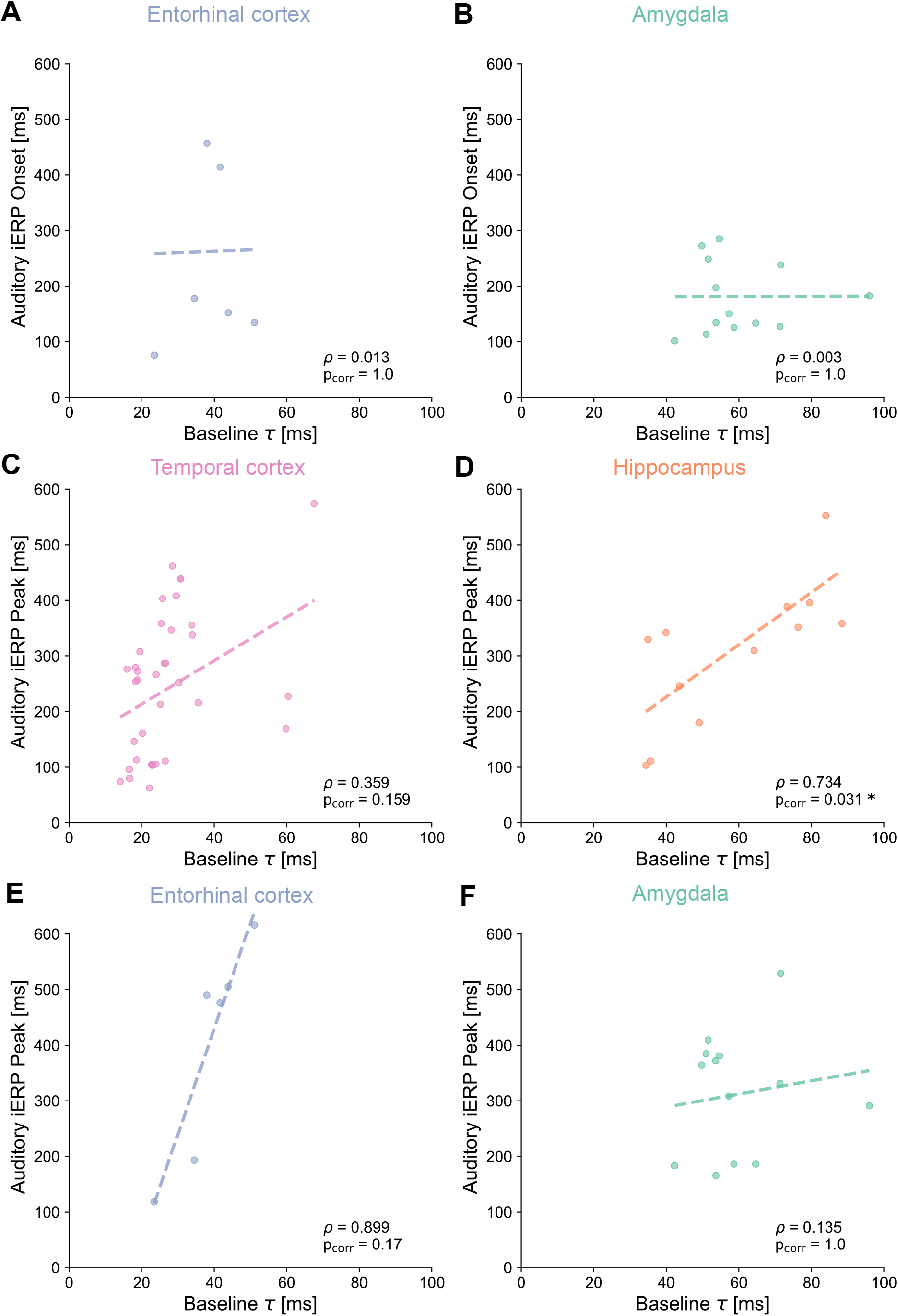
Relation between onset and peak latencies of auditory responses and intrinsic timescales at rest within brain regions. A, B Intrinsic timescales τ at baseline (x-axis) and onsets of iERPs (y-axis) for entorhinal cortex (A) and amygdala (B) are not correlated. C, D, E, F Intrinsic timescales τ at baseline (x-axis) and peaks of iERPs (y-axis) for temporal cortex (C), hippocampus (D), entorhinal cortex (E), and amygdala (F). Only the regression of intrinsic timescales on iERP peak in the hippocampus is significant. The ranges in both axes are kept constant across panels to facilitate comparison.

**Supplementary Figure 4-1.**
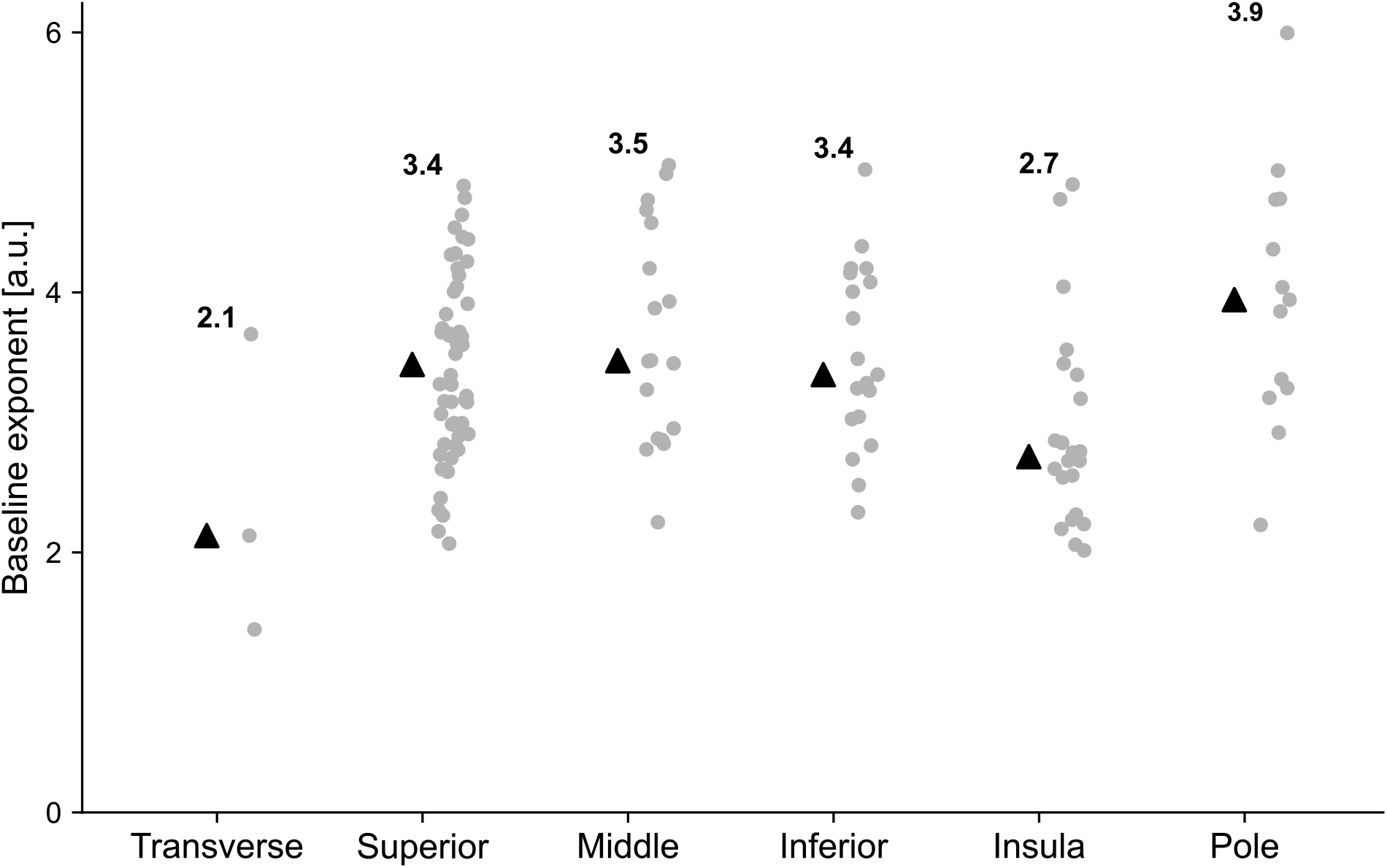
20-35 Hz spectral exponent at baseline across cortical sub-divisions. Spectral exponents of cortical electrodes, divided by anatomical subregion. Black triangles indicate median values, along with the actual value written on top. Transverse: transverse temporal gyrus; Superior: superior temporal gyrus and sulcus; Middle: middle temporal gyrus; Inferior: inferior temporal gyrus and sulcus; Insula: inferior circular sulcus of insula; Pole: temporal pole.

**Supplementary Figure 4-2.**
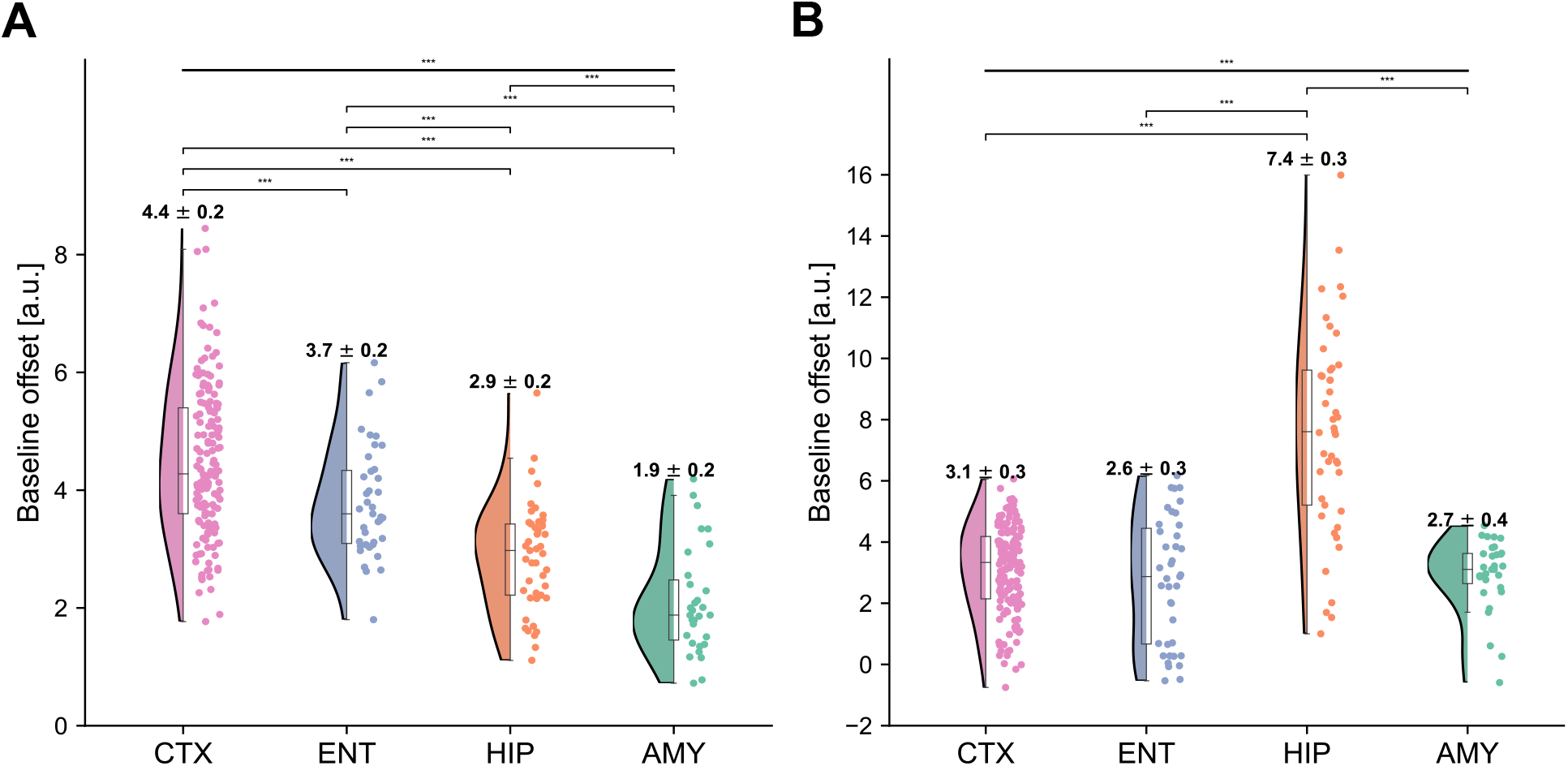
20-35 Hz spectral offset across brain regions. A. The spectral offset at 20-35 Hz is plotted for each electrode, and shows a significant main effect of region (F(3,256)=72.995, p<10^−16^), with the temporal cortex having the highest offset, followed by the entorhinal cortex, hippocampus, and amygdala. B. Spectral offset at 80-150 Hz shows a significant effect of region (F(3,256)=73.832, p<10^−16^), with the hippocampus having the highest value. The corresponding exponents can be found in Figure 4, main manuscript.

**Supplementary Figure 4-3.**
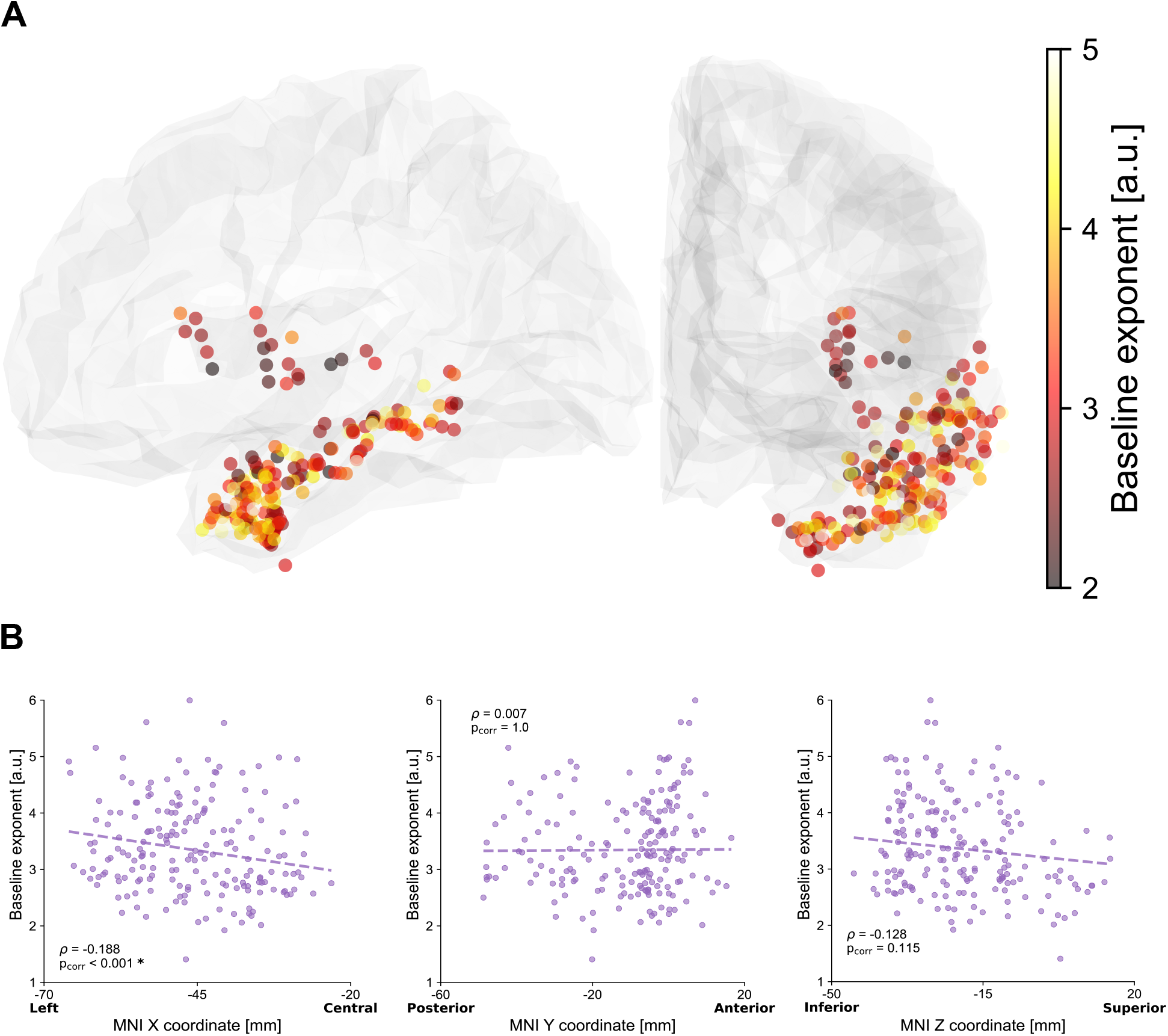
20-35 Hz spectral exponent at rest in cortical electrodes. A. Anatomical organization of spectral exponent throughout the temporal and entorhinal cortices, with the color map quantifying the exponent for each electrode. B. Correlations between MNI coordinates and spectral exponent across electrodes show significance for the X (medio-lateral) and Z (inferior-superior) directions (ρ_X_=-0.188, p_X_=9.988⨉10^−4^; ρ_Y_=0.007, p_Y_=1.0; ρ_Z_=-0.128, p_Z_=0.115; p-values after Bonferroni correction). For display purposes, all electrodes have been projected to the left hemisphere.

**Supplementary Figure 4-4.**
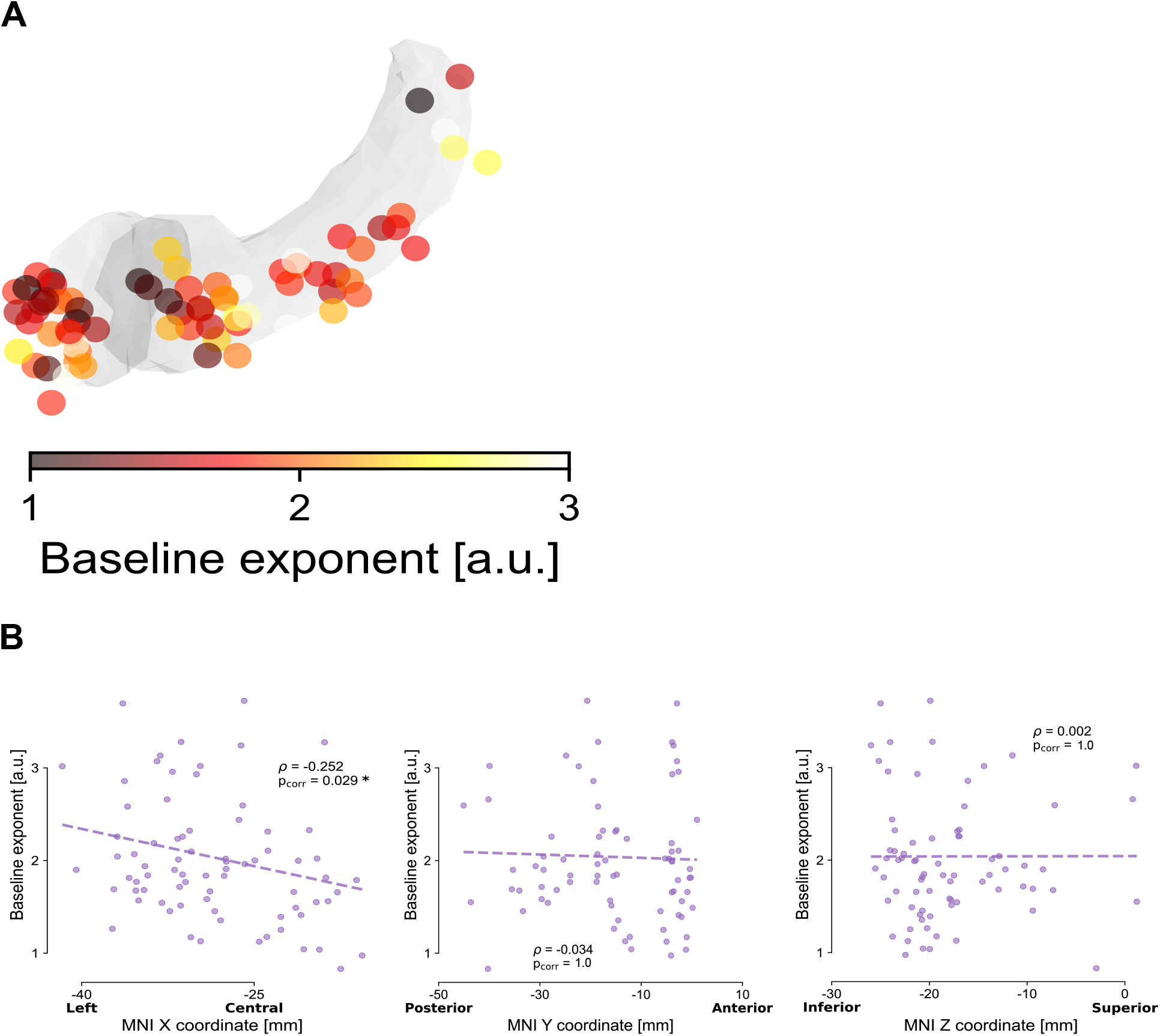
20-35 Hz spectral exponent at rest in hippocampus and amygdala. A. Anatomical organization of spectral exponent throughout the hippocampus and amygdala, with the color map quantifying the exponent for each electrode. B. Correlations between MNI coordinates and spectral exponent for each electrode show significance only for the X (left-central) direction (ρ_X_=-0.252, p_X_=0.029; ρ_Y_=-0.034, p_Y_=1.0; ρ_Z_=0.002, p_Z_=1.0; p-values after Bonferroni correction). For display purposes, all electrodes have been projected to the left hemisphere.

**Supplementary Figure 4-5.**
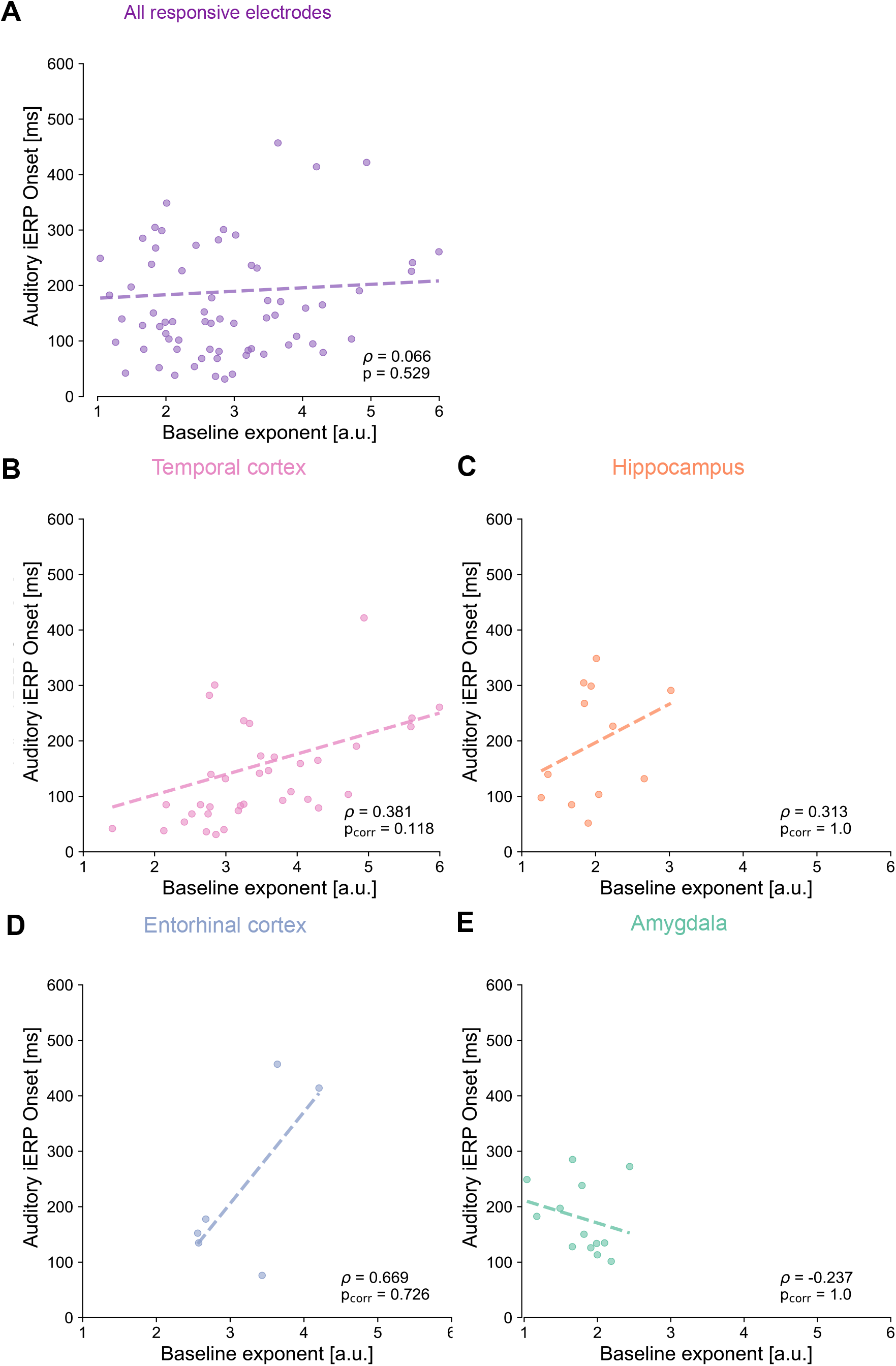
Relation between onset latency of auditory responses and 20-35 Hz spectral exponent at rest. Regression of spectral exponent at baseline (x-axis) on onsets of iERPs (y-axis) for all responsive electrodes (A), temporal cortex (B), hippocampus (C), entorhinal cortex (D), and amygdala (E). None of the regression is significant. The reported p values have been corrected for multiple comparisons across regions. Notice the ranges in both axes are kept constant.

**Supplementary Figure 4-6.**
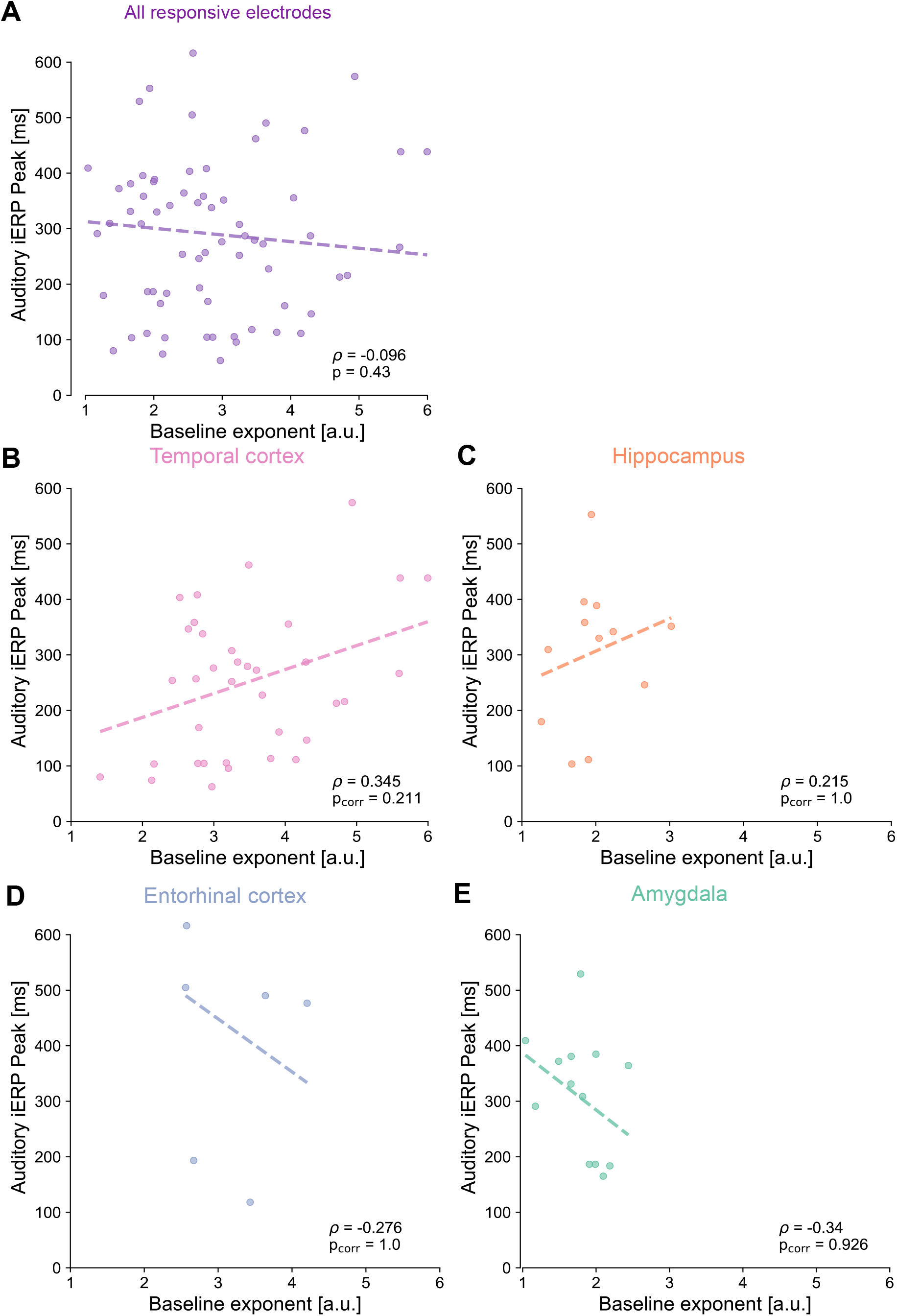
Relation between peak latency of auditory responses and 20-35 Hz spectral exponent at rest. Regression of spectral exponent at baseline (x-axis) on peaks of iERPs (y-axis) for all responsive electrodes (A), temporal cortex (B), hippocampus (C), entorhinal cortex (D), and amygdala (E). None of the regression is significant; all reported p values have been corrected for multiple comparisons. Notice the ranges in both axes are kept constant.

